# Dexamethasone induces senescence of lung epithelial cells and augments TGF-β1-mediated production of the fibrosis mediator serpin E1 (plasminogen activator inhibitor-1)

**DOI:** 10.1101/2021.11.29.470337

**Authors:** Francesca. L. Longhorne, Holly N. Wilkinson, Matthew J. Hardman, Simon P. Hart

**Affiliations:** Respiratory Research Group, Hull York Medical School, Castle Hill Hospital, Hull, United Kingdom; Centre for Atherothrombosis and Metabolic Disease, Hull York Medical School, Hull, United Kingdom

**Author notes:** Corresponding author: (SH).

## Abstract

**Background:** Idiopathic pulmonary fibrosis (IPF) is a progressive, incurable scarring disease of the lungs with a prognosis worse than most cancers. Pathologically, IPF is characterised by upregulation of the pro-fibrotic cytokine transforming growth factor-β1 (TGF-β1), activation of fibroblasts, and deposition of collagen in the alveolar interstitium. Recent evidence has highlighted the role of senescent type 2 alveolar epithelial cells in the pathogenesis of IPF. In a clinical trial, a treatment regimen containing a corticosteroid drug accelerated pulmonary fibrosis leading to more hospitalizations and deaths, particularly in patients with telomere shortening which drives cellular senescence.

**Aim:** To investigate the potential pro-fibrotic actions of corticosteroids on lung epithelial cells *in vitro*, including effects on cellular senescence and interactions with TGF-β1.

**Methods:** The synthetic glucocorticoid dexamethasone (DEX) was incubated with A549 and BEAS-2B human lung epithelial cells in the presence or absence of TGF-β1. Cellular senescence was assessed by morphology, senescence-associated beta-galactosidase (SA β-Gal) expression, and qPCR for transcription of senescence-associated molecular markers. Conditioned media were screened for growth factors and cytokines and cultured with human lung fibroblasts. An IPF lung tissue RNA array dataset was re-analysed with a focus on senescence markers.

**Results:** DEX induced senescence in lung epithelial cells associated with increased p21 (CDKN1A) expression independently of p16 (CDKN2A) or p53 (TP53). DEX amplified upregulation of the pro-fibrotic mediator serpin E1/plasminogen activator inhibitor-1 (PAI-1) in the presence of TGF-β1. The senescence-associated secretory phenotype from lung epithelial cells treated with DEX plus TGF-β1-treated contained increased concentrations of GM-CSF and IL-6 and when incubated with primary human lung fibroblasts there were trends to increased senescence and production of fibrosis markers. Upregulation of senescence markers was demonstrated by analysis of an IPF transcriptomic dataset.

**Discussion:** DEX induces senescence in lung epithelial cell lines *in vitro* and interacts with TGF-β1 to amplify production of the pro-fibrotic mediator serpin E1 (PAI-1). This may be a mechanism by which corticosteroids promote pulmonary fibrosis in susceptible individuals. Serpin E1/PAI-1 is a potential druggable target in pulmonary fibrosis.

## Introduction

Idiopathic pulmonary fibrosis (IPF) is an incurable lung disease with a median survival of only 2-5 years from diagnosis due to progressive lung scarring, loss of lung function, and respiratory failure (1). Pathologically, IPF is characterised by upregulation of pro-fibrotic cytokines including TGF-β1, fibroblast activation and accumulation, irregular deposition of collagen in the alveolar walls, and honeycomb change (2)(3). The cause of IPF remains unknown, but well-established risk factors include older age (4), prior tobacco smoking, and telomere shortening, all of which are linked with cellular senescence (5)(6)(7)(8).

Recent evidence has highlighted the role of senescent type 2 alveolar epithelial (AT2) cells in the pathogenesis of IPF (9, 10). Key senescence characteristics have been consistently observed in IPF lungs, including increased p21 (CDKN1A), p16 (p16INK4a or CDKN2A), p53 (TP53), and β-galactosidase (9)(11–13). Applying single cell RNA sequencing to IPF lung tissue, Yao et al demonstrated that AT2 alveolar epithelial cells had senescent transcriptomic characteristics and in a conditional knockout mouse model, triggered senescence of AT2 alveolar epithelial cells was sufficient to induce pulmonary fibrosis (14).

Historically, treatment of IPF with corticosteroid drugs, often in combination with other immunosuppressants, was adopted worldwide although clinical trial data were lacking (15). In 2012, interim analysis of the PANTHER-IPF trial (16) reported that combination treatment with the steroid prednisone, azathioprine, and N-acetylcysteine (NAC) caused more deaths and hospitalizations compared with the placebo group. The adverse events were predominantly respiratory, indicating that combination treatment had accelerated pulmonary fibrosis progression rather than attenuating it. A post hoc subset analysis of PANTHER showed that the adverse signal with steroid-containing therapy was driven by a subgroup of patients who had short telomeres (17) and were hence predisposed to premature cellular senescence.

We aimed to investigate whether cellular senescence played a role in the fibrosis-propagating action of corticosteroids, and their interactions with TGF-β1, using human lung epithelial cell lines *in vitro*. We report that the steroid dexamethasone (DEX) induced senescence in lung epithelial cells, synergistically increased transcription of the pro-fibrotic mediator serpin E1/plasminogen activator inhibitor-1 (PAI-1) in the presence of TGF-β1, and the senescence-associated secretory profile (SASP) produced by these cells may induce fibrosis markers and senescence in human lung fibroblasts. Published RNA-Array data (18) from IPF lung samples were re-analysed to link our *in vitro* with *in vivo* findings.

## Materials & methods

### Cell Culture and Treatments

A549 and BEAS-2B cells (ATCC, Middlesex, UK) were maintained in DMEM media (Thermo Fisher Scientific) with 1% Penicillin/Streptomycin, 1% L-glutamine and 10% foetal bovine serum (Thermo Fisher Scientific) at 37°C with 5% CO_2_. Cells were treated for 48 hours with a carrier control (media + phosphate buffered saline), dexamethasone (Hameln Pharmaceuticals) at concentrations between 10^-8^M and 10^-5^M, TGF-β1 (5 ng/mL, BioRad), or a combination of DEX (10^-6^M) and TGF-β1. Primary lung fibroblasts were grown from human lung tissue explants (with ethics committee approval, REC 12/SC/0474) and maintained in DMEM medium up to 7 passages.

### Cell Counting and Staining

Cells were lifted using Hepes-buffered saline/EDTA and resuspended in DMEM media before manual counting from a set volume. For morphology staining, cells were washed in PBS then stained with Shandon™ Kwik-Diff™ (Thermo Scientific). For senescence-associated beta-galactosidase (SA β-Gal) staining, the methodology of the chromogenic assay from Debacq-Chainiaux et al. (2009) was adapted and positive blue cells quantified. Cells were visualised on a Nikon E400 microscope using brightfield (Image Solutions, Inc. Michigan, US or SPOT imaging, Michigan, US).

### Livecyte

Cells were treated with conditions and run on the Phasefocus Livecyte™ microscope for 48 hours. Data were analysed using the Livecyte™ programme for parameters related to cell morphology, dry mass, and velocity (Phasefocus Ltd, Dashboards: User Manual, Version 3.0.1).

### Intracellular and Cell Surface Protein Expression

For surface expression, cells were blocked in 0.5% bovine serum albumin (BSA) (Fisher) and incubated with primary conjugated antibodies for E-Cadherin (CD324), N-Cadherin (CD325) or EpCAM (CD326), with corresponding isotype controls (Biolegend®, California, USA). Cells were analysed using flow cytometry (BD FACSCalibur™), until 10,000 events had been recorded.

For intracellular mTOR expression, A549 cells were fixed, blocked and washed with True-Phos™ Perm Buffer (Biolegend®, California, USA), 0.5% BSA and Cell Staining Buffer (Biolegend®, California, USA). Cells were incubated with an Anti-Hu/Mo Phospho-S6 (Ser 235, Ser 236) antibody and corresponding isotype control (eBioscience™, Invitrogen). Samples were analysed using flow cytometry as above.

### Conditioned media collection

A549 conditioned media (C/M) were generated by treating A549 cells for 48 hours, discarding supernatants, then adding fresh DMEM media collected after two days. All samples had matched protein concentrations of approximately 4μg/mL. C/M collected from treated A549 cells were screened for growth factors and cytokines using a Human Inflammation and Human Growth antibody array, screening for 40 and 41 targets (Abcam). Mediators showing differing treatment effects results on all blot replicates were re-plotted making densities relative to the control condition and positive cytokines results confirmed by ELISA. C/M were diluted 1:4 with fresh medium when added to primary lung fibroblasts over 10 days. Medium was changed every two days.

### qRT-PCR

RNA was extracted from cells using Invitrogen™ TRIzol® reagent (Thermo Fisher Scientific) and chloroform for fraction separation. The RNA fraction was used with the Invitrogen™ PureLink™ RNA Mini Kit (Thermo Fisher Scientific) as per the manufacturer’s instructions. RNA was adjusted to 1μg/10μL using NanoDrop (SimpliNano, Biochrom) then cDNA synthesised using a thermocycler (Techne, TC-412) and MultiScribe® Reverse Transcriptase from the TaqMan Reverse Transcription Reagents kit (Applied Biosystems). Gene expression was analysed by qPCR on a CFX Connect™ thermocycler (Bio-Rad, Hertfordshire, UK), using Takyon™ SYBR mastermix (Eurogentec, Hampshire, UK). Quality control measures included using amplification curves between cycles 20-35, efficiency values between 90-110% and R2 value close to 1. Any values that fell outside of these cut-offs were not used.

### Cytokine Quantification

Inflammation cytokine array blots (Abcam) were incubated with A549 C/M as per the manufacturer’s instructions. Blots were visualised using the C-DiGit® scanner system (LI-COR® Biosciences) and densities quantified using ImageJ (version 1.52t). Confirmation of positive results was performed using ELISA kits for IL-6 and GM-CSF (Abcam).

### Zymography

Acrylamide gels (10%, v/v) containing 0.2% (w/v) porcine gelatin (Thermo Fisher Scientific) were resolved with a Triton X-100 buffer, stained with 0.1% Amido black (Fisher) and washed with 10% acetic acid (Fisher Science) as previously described (19, 20). Band densities were quantified using ImageJ (version 1.52t).

### RNA-Array Data Set Analysis

A published data set from DePianto et al. (18) included 40 IPF lung samples and 8 control lung samples. Data were re-analysed using R (version 3.6.1) to compare the IPF samples to the control samples for generation of principal component analysis (PCA) plots and heatmaps of the most statistically significant variable genes. Functional annotation and GO enrichment analysis for differentially expressed genes (DEG) were carried out using the online Database for Annotation, Visualization and Integrated Discovery (DAVID) (version 6.8) (21–25).

### Statistical analysis

One-way or two-way ANOVA with Dunnett’s multiple comparison were performed. For SA β-Gal counts, data was plotted using percentages with a negative binomial regression. Analyses were performed using SPSS (version 26), GraphPad (Version 8.0; GraphPad Software, California, US) and R (Version 3.6.2; R Foundation for Statistical Computing, Vienna, Austria).

## Results

### DEX treatment does not affect EMT but leads to changes in cell number and morphology

We first investigated whether glucocorticoid treatment could induce epithelial-to-mesenchymal transition (EMT). TGF-β1 significantly reduced surface expression of epithelial markers E-Cadherin (CD324) and EpCAM (CD326) and upregulated the mesenchymal marker N-Cadherin (CD325) in A549 cells, but DEX had no effect at concentrations up to 10^-5^M either alone or in combination with TGF-β1 (S1 Appendix, a).

Incubation of lung epithelial cells with DEX led to clear differences in cell numbers and morphology as assessed by light microscopy. Cell numbers significantly decreased with increasing concentrations of DEX, and cells were enlarged compared with the carrier control, with significantly increased cell perimeter measurements (Fig 1, a-c). Similar morphological changes were seen with DEX treatment of BEAS-2B cells (S1 Appendix, b).

**Fig 1:**
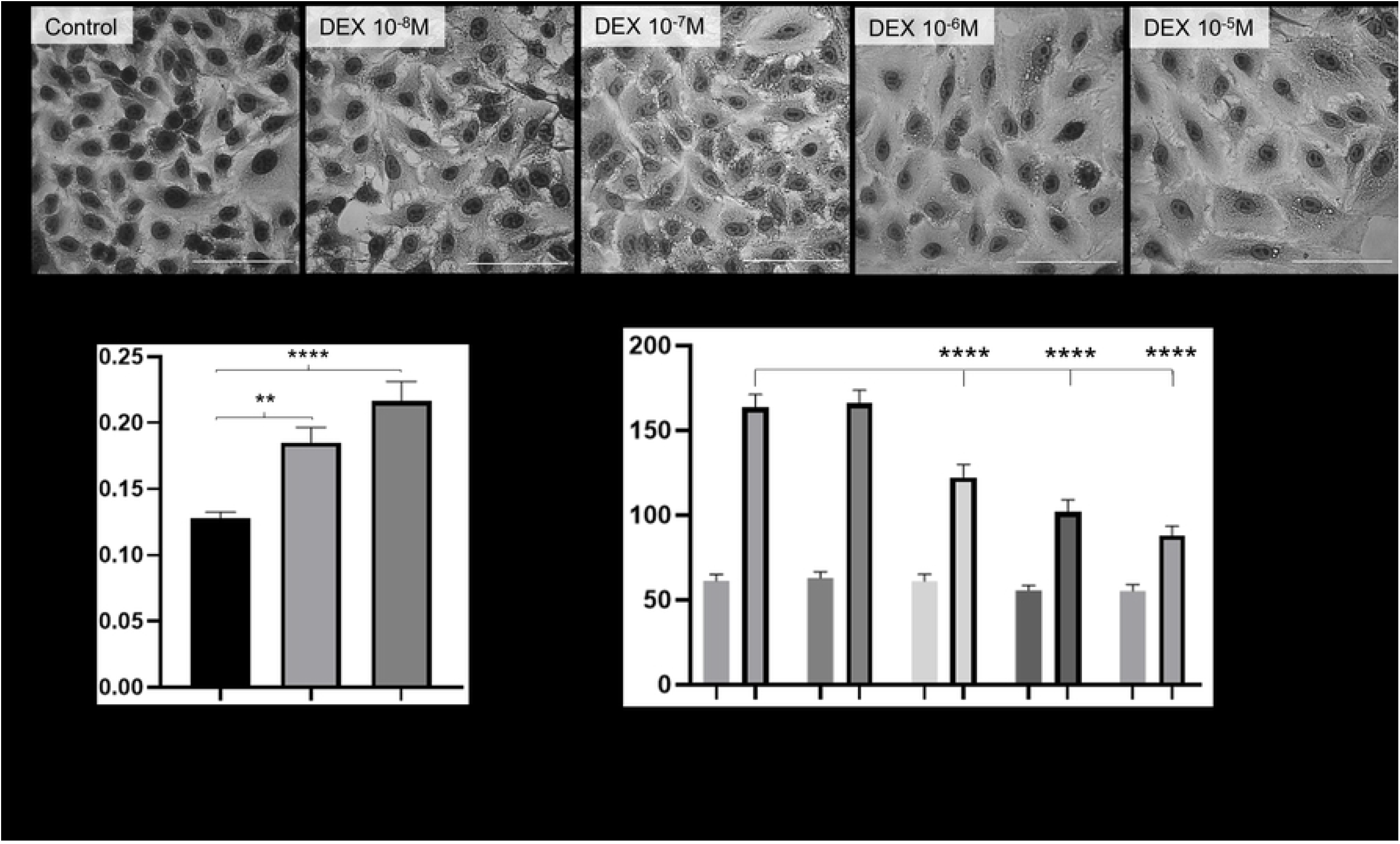
DEX induces senescence in lung epithelial cells. a) Epithelial cell size increased significantly over 48-hours, from small uniform cells to enlarged, spread cells, with higher concentrations of DEX, further shown by increased average cell perimeter sizes (b, One-way ANOVA with Dunnett’s multiple comparisons (**/****:p≤0.01/ ≤0.0001) compared to control). Scale bar 0.1mm, error bars show SEM. c) Epithelial cell counts significantly decreased over 48-hours with higher concentrations of DEX (Two-way ANOVA with Dunnett’s multiple comparisons (****: p≤0.0001) compared to control).

### DEX induces expression of senescence-associated β-galactosidase (SA β-Gal)

Cellular senescence is characterised by halted cell division and increased cell size. As further evidence of senescence following DEX treatment, there was a concentration-dependent increase in the senescence marker SA β-Gal (Fig 2, a-b). DEX-induced expression of SA β-Gal was also seen in BEAS-2B cells (S1 Appendix, c). The combination of the observed changes in A549 cell morphology with positive SA β-Gal staining indicated that DEX was inducing cellular senescence.

**Fig 2:**
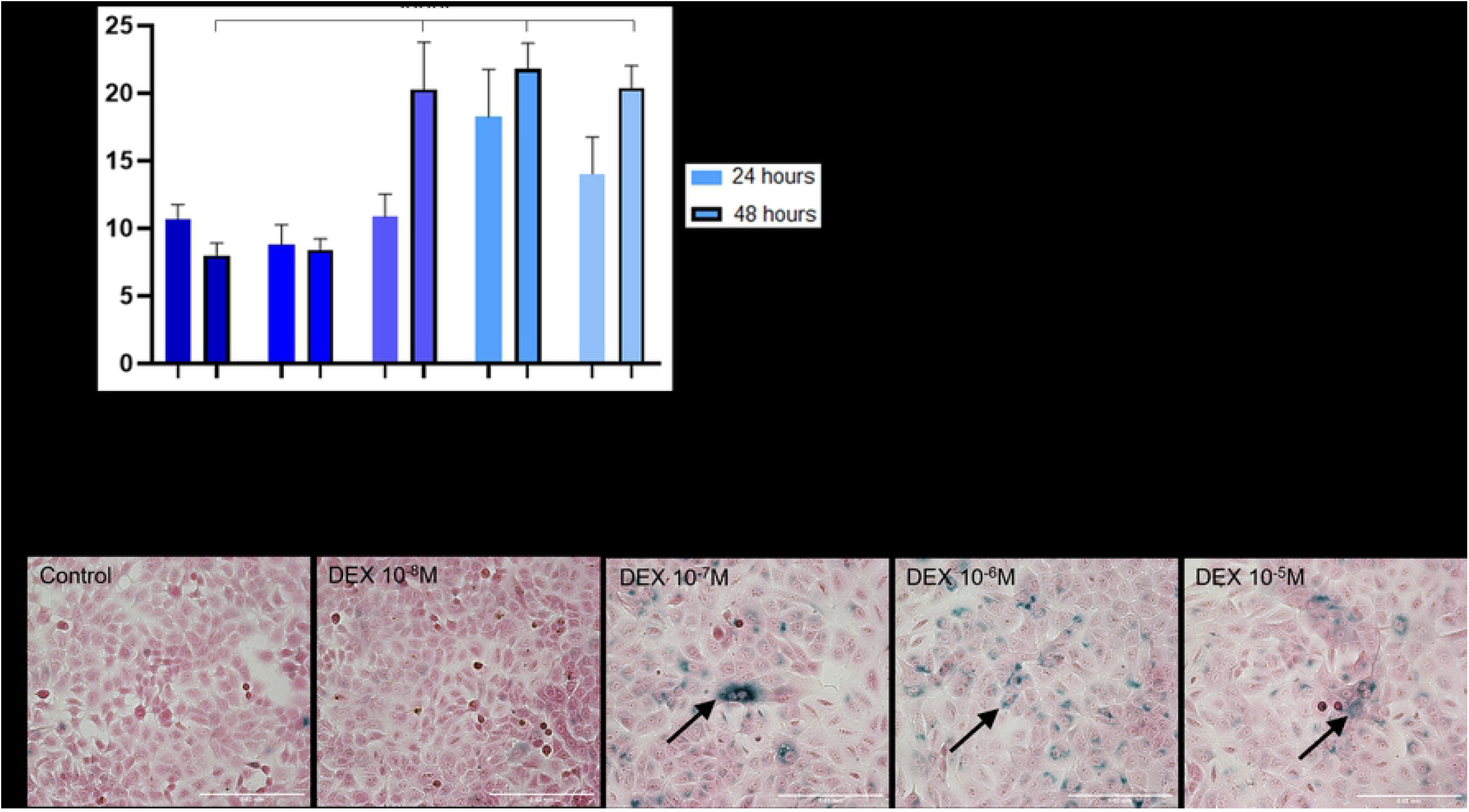
DEX induces senescence in lung epithelial cells. To assess senescence, SA β-gal staining was used. The number of positive blue cells (black arrows) significantly increased over time (a) and with stronger DEX concentrations (b, DEX 10^-7^M, 10^-6^M, 0^-5^M, negative binomial regression analysis, ****: p≤0.0001) compared to the carrier control and weakest DEX centration (10^-8^M). Scale bar 0.02mm, error bars show SEM.

### DEX-induced senescence is associated with transcription of p21 and Serpin E1/PAI-1 but not p16 or p53

Gene expression of senescence markers in lung epithelial cells was investigated after incubation DEX with or without TGF-β1, a potent inducer of fibrosis (Fig 3, a). DEX alone significantly increased transcription of p21 (*CDKN2A*) in A549 cells at 24 and 48 hours, and in BEAS-2B cells at 48 hours (Fig 3, b), in keeping with the time course of induction of senescence. In contrast, DEX-induced senescence was not associated with increased transcription of p16 (*CDKN1A*), p53 (*TP53*), or Serpin E1/PAI-1 (*SERPINE1*). However, in the presence of TGF-β1, DEX synergistically enhanced the increase in Serpin E1/PAI-1, an effect that was seen in both A549 and BEAS-2B cells (Fig 3, a-b).

**Fig 3:**
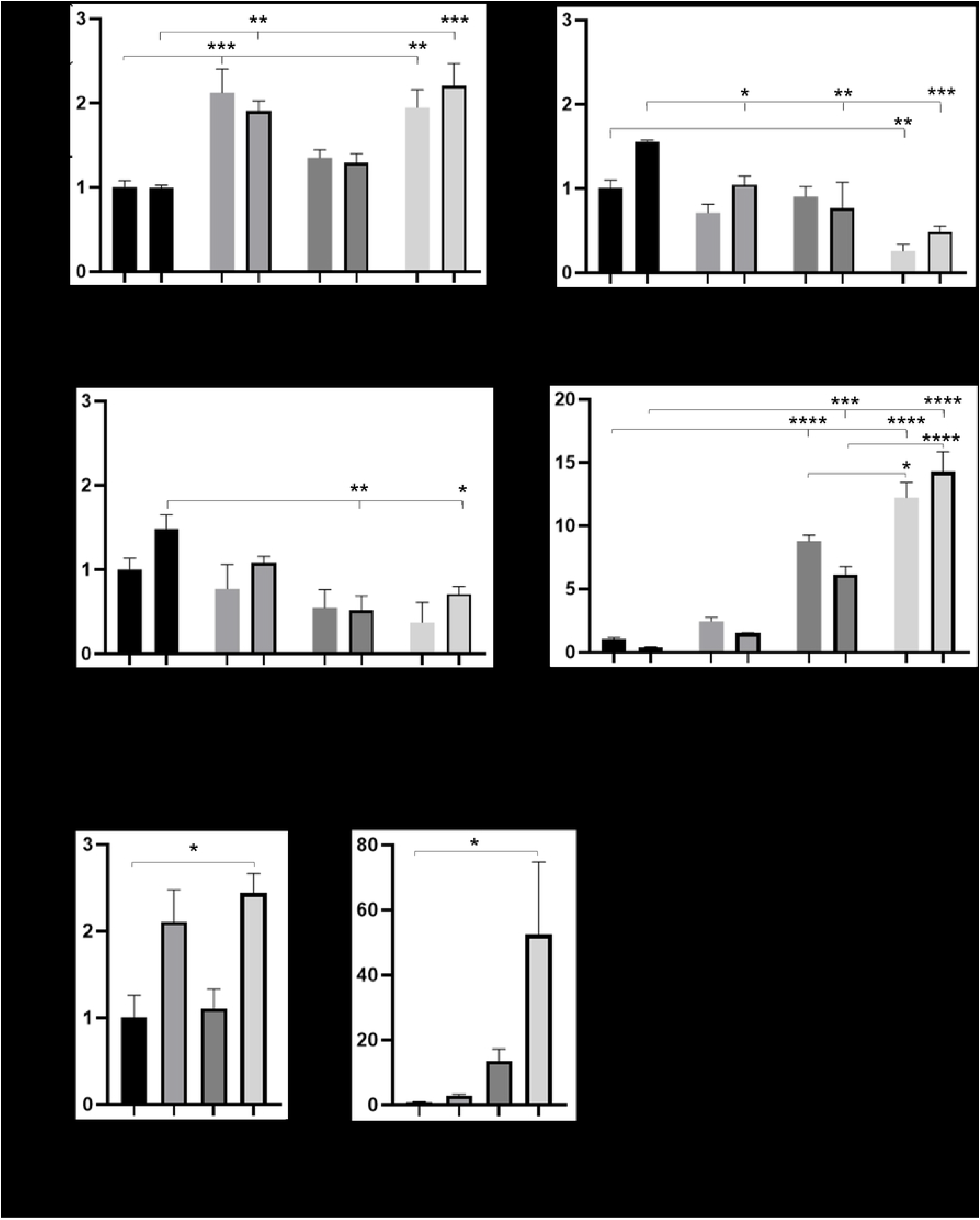
Senescence associated genes in lung epithelial cells. a) DEX treatments caused significant increases in the senescence marker *CDKN2A* (p21), and TGF-β_1_ treatments caused significant increases in the senescence and fibrosis marker *SERPINE1* (Serpin E1 or PAI-1) in alveolar epithelial (A549) cells. Interestingly, a combination of TGF-β_1_ and DEX caused a significant additive effect in *SERPINE1* expression Significant decreases were seen in *TP53* (p53) and *CDKN1A* v(p16) following treatments mainly at 48 hours. b) Similar patterns were seen in bronchial epithelial (BEAS-2B) cells at 48 hours; combination treatments caused significant increases in the marker *CDKN2A* (p21) and *SERPINE1* (serpinE1/PAI-1). Gene expression displayed as a ratio to the control treatment, n=3, error bars denote SEM. Outlined bar denotes 48h (a only). Two-way ANOVA with Dunnett’s multiple comparison or Sidak’s multiple comparison. */**/***/****:p≤0.05/ ≤0.01/ ≤0.001/ ≤0.0001.

### DEX-induced senescence is not associated with mTOR activation

Summer et al. (26) reported that the mTOR axis was increased in senescent lung epithelial cells and that when mTOR was blocked, cellular senescence was reduced. However, in the presence or absence of TGF-β1, DEX-induced senescence in A549 cells was not associated with any demonstrable changes in rapamycin-inhibitable mTORC1 activation as assessed by pS6RP phosphorylation (S2 Appendix).

### Conditioned media components

Conditioned media (C/M) from DEX-treated A549 cells were collected to analyse potential components of a senescence-associated secretory phenotype (SASP). C/M collected from treated A549 cells were screened for growth and inflammation factors using multiplex array blots (Fig 4, a) and positive results confirmed by ELISA (Fig 4, b). Concentrations of growth factors were universally low in A549 conditioned media with all treatments. In terms of inflammatory mediators, the most varied cytokines in the multiplex array blots (Fig 4, a) were granulocyte-macrophage colony-stimulating factor (GM-CSF) and interleukin-6 (IL-6). ELISA confirmed that in response to TGF-β1, IL-6 production increased significantly. DEX alone suppressed IL-6 as expected, but in the presence of TGF-β1 the increase in IL-6 was resistant to suppression by DEX (Fig 4, b). GM-CSF was suppressed by DEX, TGF-β1 alone had no effect, but the combination of DEX + TGF-β1 appeared to increase GM-CSF although this could not be confirmed statistically (Fig 4, b). The concentrations of IL-6 and GM-CSF fell within pathologically meaningful ranges reported in the literature (27–31).

**Fig 4:**
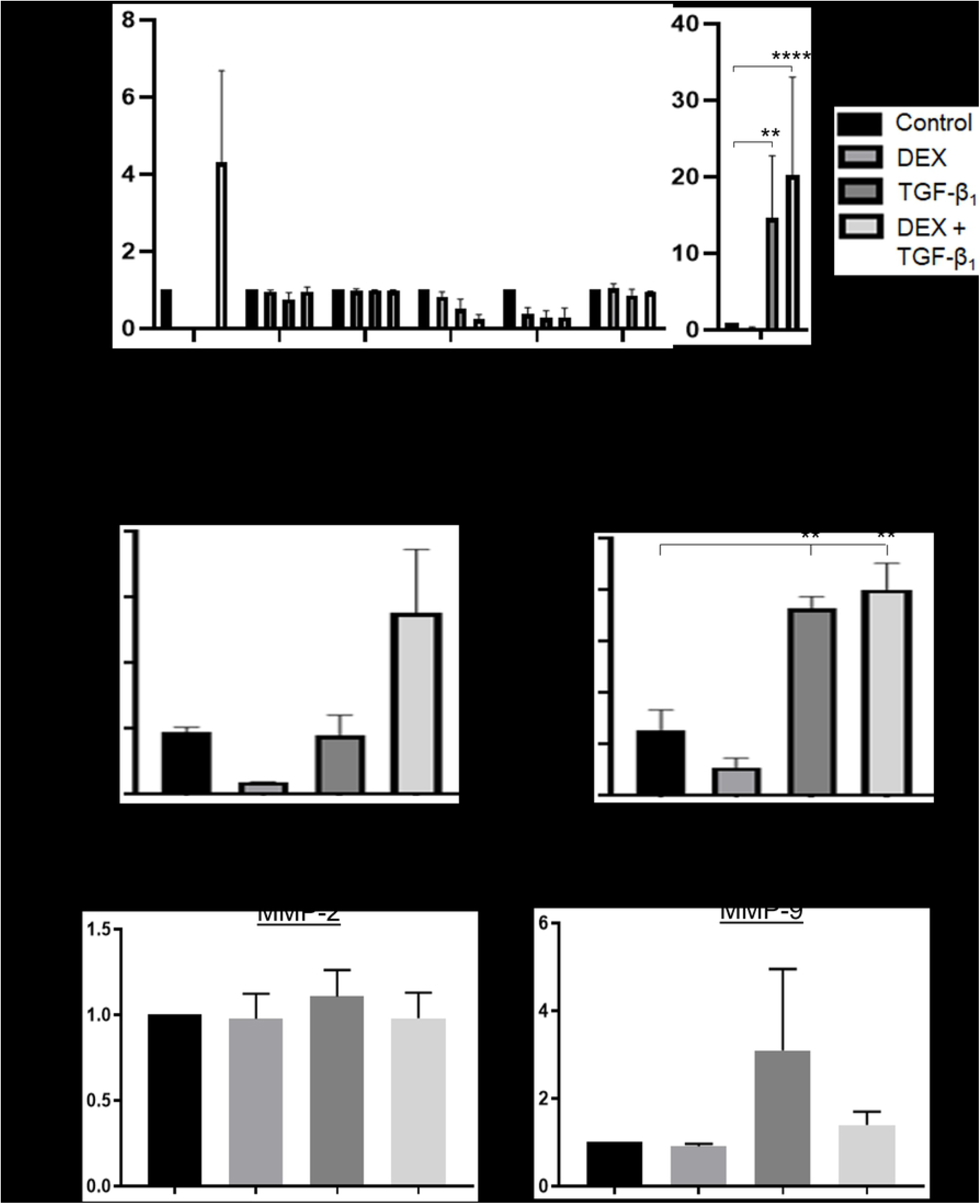
SASP components in lung epithelial cells. Measuring SASP on cytokine array blots (a) revealed two specific cytokine increases, GM-CSF with the dual condition and IL-6 with the TGF-β_1_ conditions, which were further quantified and confirmed by ELISA (b). Error bars denote SEM. Two-way ANOVA with Dunnett’s multiple comparison for cytokine array data and One-way ANOVA with Dunnett’s multiple comparison for ELISA data. **/**** p≤0.01/ ≤0.0001. Quantification of MMPs in SASP (c) revealed no change in MMP-2 and variable results of MMP-9. Error bars denote SEM with no statistical significance (One-way ANOVA).

### No evidence of MMPs in conditioned media from DEX-treated lung epithelial cells

Matrix metalloproteinases (MMPs) are able to degrade and remodel ECM proteins including collagens, and defective production or neutralization of MMPs may contribute to fibrosis (32, 33). Two MMPS of particular interest are MMP-2 and MMP-9 which are upregulated in the fibrotic lung (34, 35). Zymography was used to look for activity of MMPs in C/M from A549 cells (Fig 4, c). There was no difference in MMP-2 activity between conditions. For MMP-9, activity increased with TGF-β1 but DEX had no effect.

### Effects of conditioned media from DEX-treated A549 cells on primary lung fibroblasts

To determine potential pro-fibrotic effects of the SASP, we cultured human lung fibroblasts in C/M from senescent lung epithelial cells. After 10 days’ incubation with C/M from DEX-treated A549 cells, primary human lung fibroblasts lost their elongated, spindle-like form and became enlarged and more rounded, with more SA β-Gal positive cells indicating senescence (S3 Appendix, a). The morphology of fibroblasts made quantification challenging and statistical significance could not be demonstrated. Fibroblasts incubated with C/M from TGF-β1-treated A549 cells looked similar to the control cells in both morphology and staining. The dual condition cells demonstrated some spindle-like cells mixed with enlarged, blue positive cells.

Fibrosis- and senescence-associated gene transcription by primary human lung fibroblasts after 10 days of C/M incubation was also explored (S3 Appendix, b). After incubation with C/M from DEX-treated A549 cells, there were trends to increased expression of the fibrosis-associated genes Collagen 1, Collagen 3, Fibronectin, and Vimentin. Similar levels were seen in the dual C/M cells, indicating that the DEX-induced A549 C/M had a dominant effect on fibroblast gene expression over TGF-β1. Overall, TGF-β1 C/M had the same effect as the carrier control. There was no detectable induction of senescence-associated genes p21 or Serpin E1 in fibroblasts.

### IPF patient RNA-Array lung profiles: linking *in vitro* to *in vivo*

Re-analysis of a bulk transcriptomic dataset (18) of IPF lung samples (29 explants, 11 biopsies) confirmed that IPF lung samples were clustered away from control lung samples, with two distinct populations on the PCA plot and the heatmap of the 250 most variable genes (Fig 5, a). 14 cellular processes involving the most differentially expressed genes were highlighted using DAVID (Database for Annotation, Visualization and Integrated Discovery), including upregulation of ‘ECM’, ‘Immune Response’ and ‘p53 signalling pathway’ and downregulation of ‘cell adhesion’ and ‘angiogenesis’ (Fig 5, b). Enrichment analysis of the 10 upregulated pathways showed the fold change of the genes in those pathways, with ‘Tissue inhibitors of matrix metalloproteinases (TIMP)’ and ‘Transforming Growth Factor-beta’ being the most upregulated.

**Fig 5:**
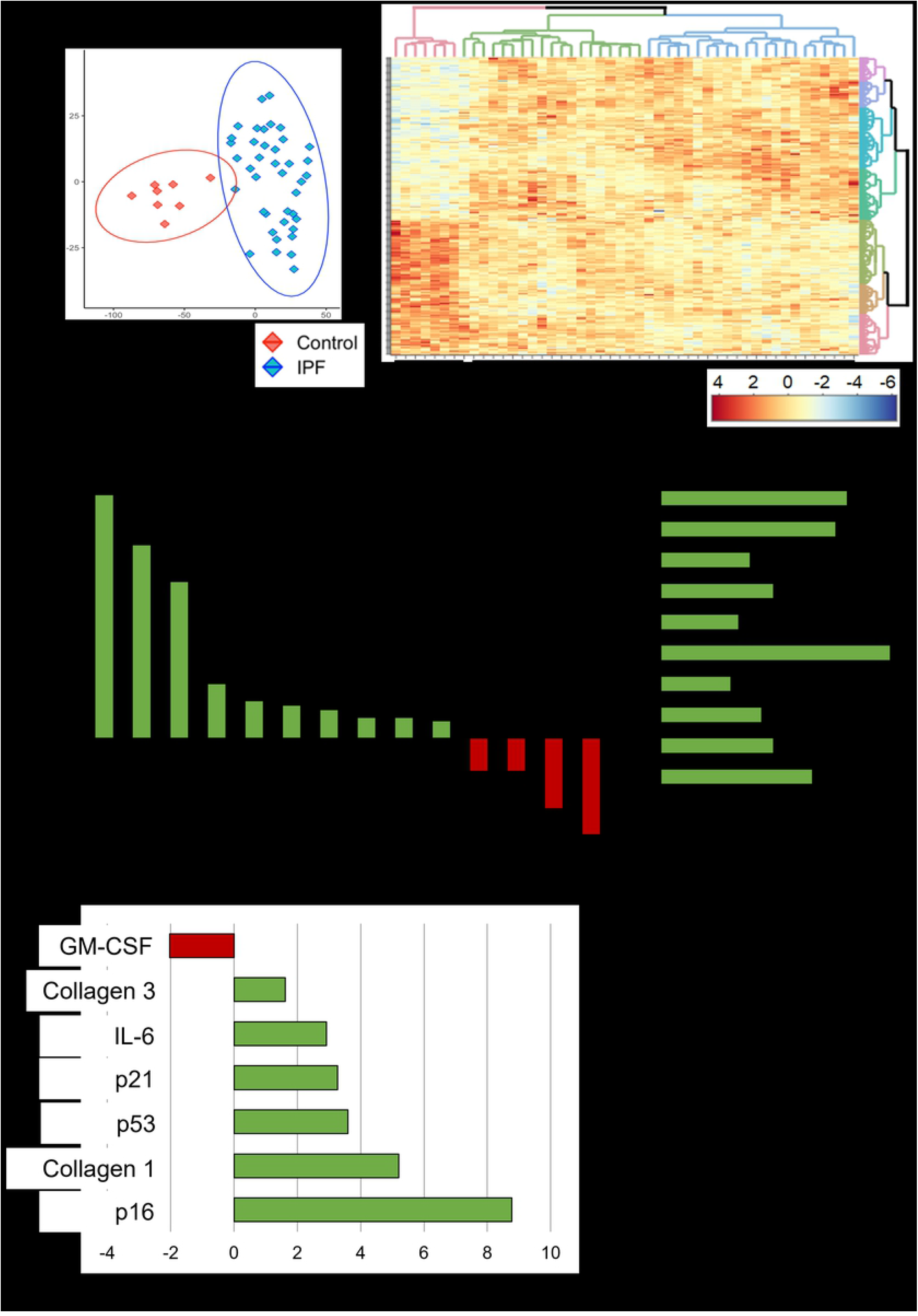
IPF patient lung profile. Published RNA-Array data analysis comparing IPF to control lung samples, shows clear sample clustering on a PCA plot and the spread of data on a heat map of 250 most variable (up- or down-regulated) genes (a). b) DAVID analysis shows the biological processes with the most up- or down-regulated genes in the IPF data set, highlighting fibrosis-related or senescence-related pathways. The enrichment graph shows the fold change of the most up-regulated genes in the IPF data (Enrichment key: I: Extracellular Matrix, II: Immune Response, III: Cell Adhesion, IV: Apoptosis, V: Tissue inhibitors of matrix metalloproteinases (TIMP), VI: Response to Drug, VII: p53 Signaling Pathway, VIII: IGF Binding Protein (IGFBP), IX: Cellular Response to FGF, X: Transforming Growth Factor-beta. Differentially expressed gene (DEG) analysis (c) of seven genes related to this body of work, shows key senescence genes are upregulated in IPF, as well as collagens and the cytokine IL-6.

Next, Gene Ontology Go enrichment analysis was used to explore genes of interest in our study: Serpin E1, p21, p16, p53, GM-CSF, IL-6 and collagens 1 and 3. Serpin E1 was not identified in the 250 most differentially expressed genes. In the IPF lung dataset, GM-CSF was downregulated while p21, p16, p53, IL-6, collagen 1 and collagen 3 were upregulated (Fig 5, c).

## Discussion

Corticosteroids exert multiple mechanisms to dampen inflammation directly or indirectly (36), particularly by repressing transcription of pro-inflammatory cytokines (37). Anti-inflammatory treatment with corticosteroids was widely used for IPF on the basis that fibrosis ensued because of chronic inflammation, until practice changed following publication of the major adverse outcomes of the PANTHER-IPF trial in 2012. By that time, alternative concepts of IPF pathogenesis had already been proposed based whereby fibrosis occurred due to repeated injuries to the lung epithelium followed by abnormal wound repair (1, 38–40). Recently, evidence has highlighted a key role for senescence of alveolar type 2 epithelial (AT2) cells as a driver of lung fibrosis (14).

Cellular senescence, first described in the 1960s (41), is a state in which cells remain metabolically active but undergo no cell growth or death as they are in irreversible cell cycle arrest (42–44). These cells can secrete a senescence-associated secretory phenotype (SASP) which can lead to further senescence or pro-fibrotic effects (45). Senescence is thought to have beneficial anti-tumour and anti-viral effects (46–48). Senescence occurs naturally with increasing age along with reduction in telomere length and is classed as a hallmark of aging (5). Other triggers of cell senescence include oxidative stress and DNA damage caused by drugs or cytokines (5, 44, 49, 50).

Senescent cells show characteristic morphology, becoming enlarged and flatter, which we observed in A549 cells treated with pharmacologically relevant concentrations of DEX (51, 52). Peak plasma concentrations around 0.2μM (2×10^-7^M) are found in response to treatment with 6mg or oral dexamethasone daily (53), which equates to doses of prednisone used in PANTHER. We confirmed increased expression of intracellular SA β-Gal, a widely accepted biomarker for senescence (52) in response to DEX (54, 55). In the present study, p21 and Serpin E1 increased in association with DEX-induced senescence, but there was no increase in p16 or p53. Generally, cellular senescence can be induced by through one of two major mechanisms: activation of p53–p21 or p16INK4a–pRB pathways. Lehmann et al. (13) demonstrated increased p16 and p21 in IPF lung tissue versus control tissue, as well as bleomycin-induced fibrotic mouse lungs. The p53-p21 pathway has been linked to telomere-initiated cellular senescence when dysfunctional telomere shortening triggers a DNA damage response within the cell. Telomere shortening, with or without mutations in the TERT telomerase complex is a recognized risk factor for IPF. In the present study, DEX induced senescence in lung epithelial cells was mediated by p21 independently of p53 (56).

Serpin E1/PAI-1 increased in response to TGF-β1 (57), an effect with was enhanced in combination with DEX. SerpinE1/PAI-1 is produced by AT2 cells and may promote pulmonary fibrosis by inhibiting tPA/uPA and hence blocking degradation of ECM leading to its accumulation (57), and potentially and via interaction with its receptor LRP1 on fibroblasts or macrophages (58). We also recognise serpin E1/PAI-1 expression as a marker of senescence in AT2 and other cells (59–61). Serpin E1/PAI-1 can be activated by p53 (62, 63), but not in our model of p53-independent senescence induced by DEX. TGF-β1 is an archetypal inducer of tissue fibrosis by activating fibroblasts to become collagen-synthesizing myofibroblasts, and also induces epithelial cells to adopt a mesenchymal phenotype (epithelial-to-mesenchymal transition (EMT)) (64–66). Rana et al. (61) showed that TGF-β1 induced senescence in primary rodent ATII cells *ex vivo* associated with increased serpin E1/PAI-1 production, and that blocking PAI-1 diminished senescence and SASP-mediated activation of alveolar macrophages.

The senescence associated secretory phenotype (SASP) describes a mixture of growth factors and cytokines released by senescent cells that can have impactful effects on nearby cells (6, 67–70). In this study, we analysed conditioned media from DEX-treated senescent A549 cells with or without TGF-β1 for growth factors and cytokines that may influence fibrosis in the lung. We did not find detectable levels of growth factors implicated in fibrosis such as TGF-β, PDGF, FGF, or VEGF. DEX is a general repressor of inflammatory cytokine transcription, in keeping with our finding of suppression of several cytokines in our array. We did not find any cytokines that were upregulated is the SASP from DEX-induced senescent epithelial cells. However, combined incubation with DEX plus TGF-β, designed to mimic the milieu in the fibrotic lung, induced secretion of IL-6 and GM-CSF.

IL-6 is multifunctional inflammatory cytokine (49, 71) that has been linked to pulmonary fibrosis (72–75). In response to DEX alone, basal IL-6 production from A549 cells was suppressed as expected. As TGF-β1 is a known inducer of fibrosis, it was interesting to see IL-6 significantly increased in response to TGF-β1 at that TGF-β1-stimulated IL-6 was resistant to suppression by DEX. This finding may partly explain why corticosteroid therapy does not work for patients with IPF when there is a TGF-β1-rich milieu within the fibrotic lung. GM-CSF, involved with activation and proliferation of macrophages and immune cells (76) appeared to increase with the combination of DEX plus TGF-β1 but this result should be confirmed in further experiments.

Proteinases are important in fibrotic lung disease as they contribute to the degradation of ECM, which can aid remodelling and may be potential targets for IPF therapies. MMP-2 and MMP-9, gelatinase A and B respectively, have been shown to be increased in IPF and other chronic remodelling diseases (35, 77, 78). In this study, DEX did not affect MMP-2 or −9 from A549 cells, but TGF-β1 increased MMP-9, which itself can activate latent TGF-β1 (77).

Activated fibroblasts are the primary source of the excessive collagen-rich ECM seen in pulmonary fibrosis (79, 80). If factors within the SASP from DEX-treated epithelial cells could stimulate fibroblasts to produce more ECM, this may worsen lung fibrosis. C/M from DEX-treated A549 cells caused more positive SA β-Gal staining in primary lung fibroblasts and there were trends to increased expression of collagen and ECM-associated genes. This suggests that the effects of DEX on epithelial cells will in turn affect lung fibroblasts to become pro-fibrotic, and has the potential to effect other cells too, warranting further investigation. Whilst our RNA transcription data indicate that the SASP from DEX plus TGF-β1-conditioned A549 cells is likely to contain increased amounts of serpin E1/PAI-1, apparent contradictory effects of serpin E1/PAI-1 on fibroblasts compared with epithelial cells have been reported (81).

To link our *in vitro* data to *in vivo* evidence, we re-analysed a published bulk RNA array dataset that compared IPF lung tissue (from biopsies or explants) with control lung tissue (18). We replicated distinct transcriptomic populations of control and IPF samples (82–84). Using DAVID and enrichment analysis, we identified upregulation of senescence-associated pathways, as well as expected fibrosis pathways such as ECM. Using DEG analysis, all three senescence markers p16, p21, and p53 were increased in IPF lung. IL-6 was increased while GM-CSF decreased. Serpin E1/PAI-1 was not listed among the significantly differentially expressed genes. Fibrosis related genes collagen 1 and 3 were also upregulated. DePianto et al. (18) used whole lung tissue in their RNA array, so upregulation in epithelial cells could be masked by other cell types in the tissue. We do not know how many of the IPF subjects were taking corticosteroid therapy at the time of their biopsy or transplant.

Our study has several limitations. The use of A549 cells to model lung epithelium has been criticized due to these cells being derived from a patient with lung adenocarcinoma. In studies of DNA damage, A549 cells may exhibit heightened increased oxidative stress responses due to a mutation in KEAP1 (85). However, A549s share many features of AT2 cells (86), dysfunction of which has been heavily implicated in pulmonary fibrosis. In our model they reproducibly underwent senescence in response to pharmacologically relevant concentrations of dexamethasone, and we suggest they may be a good model to study potentially pro-fibrotic mechanisms in lung epithelium. Importantly, we reproduced our findings in BEAS-2B cells, which are ontogenically differently from A549 cells, being derived from the bronchial epithelium of a subject without cancer, transformed *in vitro* by transfection with an adenovirus 12-SV40 hybrid virus. An important caveat is the study of p16 in senescence, since A549 cells (and many other cell lines) harbour a genetic deletion in p16 that impairs the function of the p16INK4A protein (87). This is consistent with our results where we detected small amounts of p16 gene expression, but no upregulation with induction of senescence.

The experiments using SASP-containing conditioned media are constrained by the need to dilute the C/M in growth media when treating fibroblasts, meaning that any active SASP component was diluted fourfold which may diminish the ability to detect meaningful downstream effects. The variability seen, likely compounded using primary cells from different donors, prevented demonstration of statistical significance so these finding are regarded as exploratory only.

We did not study primary human AT2 cells which are challenging to isolate and short-lived in culture. We would not expect primary epithelial cells from healthy individuals to exhibit senescence in response to dexamethasone. Corticosteroids have been widely used to treat asthma for decades, both via inhaled and systemic routes (for acute exacerbations), and to our knowledge senescence of epithelial cells has never been reported in steroid-treated asthmatics. Indeed, rather than inducing senescence, there has been a single report that steroids may induce bronchial epithelial cell programmed cell death (apoptosis) (88), although the clinical relevance of these data have been challenged (89). We hypothesize that primary AT2 cells from older individuals with IPF, particularly those with short telomeres, would be more susceptible to steroid-triggered senescence. To confirm whether accelerated senescence of susceptible AT2 cells could be responsible for steroid-induced accelerated fibrosis in IPF, further research would need to study AT2 cells from lungs of IPF patients with short telomeres, who we hypothesize are predisposed to steroid-induced senescence.

In conclusion, our findings support a potential profibrotic effect of corticosteroids in lung fibrosis by inducing senescence of ‘susceptible’ AT2 cells and upregulating serpinE1 in the presence of TGF-β1. There has been interest in “senolytic” drugs that target senescent cells to die by apoptosis (90, 91), with a feasibility study in 14 IPF patients with the combination therapy of dasatinib and quercetin (91). Caution is required in attributing effects to removing senescent cells because of possible ‘off-target’ non-senolytic effects of these drugs. An alternative approach of blocking senescence is not a desirable therapeutic aim for patients due to the important role of senescence in limiting carcinogenesis and viral infections. More specific (“senomorphic”) targeting of serpin E1/PAI-1 and other senescence-associated pathways that promote fibrosis should be sought (60).

## Abbreviations

C/M: Conditioned media
DEX: Dexamethasone
EMT: Epithelial-to-mesenchymal transition
IPF: Idiopathic Pulmonary Fibrosis
NAC: N-Acetylcysteine
PAI-1: plasminogen activator inhibitor-1
SA β-Gal: senescence-associated beta-galactosidase
SASP: Senescence-associated secretory phenotype
TGF-β1: Transforming Growth factor Beta one

